# Cellular Morphology and Motility Defects are Conserved Phenotypes of Trisomy 21 Despite Heterogeneous Adhesion Mechanisms

**DOI:** 10.64898/2026.08.03.742622

**Authors:** Katelyn Rygel, Kareem Chambers, Brian Yoon, Manasi Agrawal, Samantha A. Bailey, Mitesh Patel, Cayla Wolfe, Anna Montazzoli, Taylor Bumbledare, Paige Cassidy Malcolm, Shelby Kelemen, Connor Neifert, Jordan Headen, Leah J. Kershner, Nikita Kirkise, Kristy Welshhans

## Abstract

Down syndrome is a neurodevelopmental disorder caused by the trisomy of human chromosome 21 (T21). Down syndrome is associated with a wide range of variable clinical features, including congenital heart defects and slow wound healing; however, intellectual disability is ubiquitous and results, in part, from altered neuronal connectivity. Here, we used three sets of control and T21 human fibroblasts and one set of human induced pluripotent stem cell (hiPSC)-derived cortical neurons to examine whether changes in cellular morphology and motility are consistent across cell types in Down syndrome and to elucidate the underlying mechanisms. We found that fibroblast morphology is dysregulated in all T21 fibroblast lines. Using a transwell migration assay, cellular migration is decreased in two of the three T21 fibroblast lines. T21 hiPSC-derived cortical neurons also exhibit morphological defects, including a decrease in the length of the longest neurite and growth cone area. Because of these significant changes in morphology and motility in T21 cells, we examined proteins in the focal adhesion complex, which links the intracellular cytoskeleton to the extracellular matrix and directly controls these processes. Multiple proteins in the adhesion complex, including paxillin, vinculin, talin, and RACK1, are dysregulated in T21 fibroblasts and hiPSC-derived neurons, but these changes have high inter-individual variability. Taken together, these findings suggest that altered cellular morphology and motility are conserved features of Down syndrome that arise through heterogeneous alterations in adhesion networks. Thus, this work significantly contributes to the recent literature highlighting the need for personalized medicine in Down syndrome.

## INTRODUCTION

Down syndrome is a complex developmental disorder resulting from the trisomy of human chromosome 21 (T21). It affects about 1 in every 800 live births in the United States and is the most common genetic disorder leading to intellectual disability (Antonarakis et al., 2020; de Graaf et al., 2015; Russo et al., 2024). Humans with Down syndrome and some mouse models exhibit decreased brain size (Schmidt-Sidor et al., 1990), a reduction in neuronal proliferation (Contestabile et al., 2007), and changes in neural connectivity (Kurt et al., 2004). Down syndrome can also result in a range of other phenotypes, including slow wound healing, congenital heart defects, and distinct facial and limb features (Antonarakis et al., 2020; Bartesaghi et al., 2022; Benvenuti et al., 2025; Marentette et al., 2021; Mollo et al., 2023).

Cellular morphology and motility are key processes that likely contribute to many Down syndrome phenotypes; for example, neural crest migration underlies craniofacial development and endothelial cell migration is needed for heart development. However, much about the specific cellular and molecular mechanisms underlying these phenotypes remains unknown.

Fibroblasts are mesenchymal cells involved in many cellular processes, including extracellular matrix (ECM) formation and remodeling, tissue creation and maintenance, wound repair, and growth factor secretion (Plikus et al., 2021). Thus, motility is necessary for their typical functioning. Cellular motility is largely regulated by focal adhesions, which are multiprotein, integrin-dependent adhesion sites that link the ECM to the intracellular actin cytoskeleton (Ciobanasu et al., 2012). Over 150 proteins are associated with focal adhesions, including paxillin, vinculin, talin, and receptor for activated C kinase 1 (RACK1) (Kuo et al., 2012). Focal adhesions are dynamic structures that form and turn over to regulate the speed and direction of cell motility (Gupton and Waterman-Storer, 2006; Legerstee and Houtsmuller, 2021).

Point contacts, adhesion sites in neuronal growth cones, are similar to focal adhesions in non-neuronal cells (Huttelmaier et al., 2005; Mogilner and Keren, 2009; Welshhans and Bassell, 2011). During neuronal development, neurons extend axons with growth cones at their distal tips, which sense and respond to extracellular signals to pathfind to their synaptic targets (Jung et al., 2012; Lowery and Van Vactor, 2009; Marsick et al., 2010). Point contacts regulate the speed and directionality of growth cone motility but are smaller, contain fewer proteins, and turn over faster than focal adhesions. However, many focal adhesion core proteins are also in point contacts, including paxillin, vinculin, talin, and RACK1 (Woo and Gomez, 2006). Interestingly, the composition of proteins at point contacts is more variable and dynamic than focal adhesions (Arregui et al., 1994; Renaudin et al., 1999; Robles and Gomez, 2006; Woo and Gomez, 2006).

Recruitment of adhesion proteins is a necessary process for growth cone turning and cell motility (Kerstein et al., 2017; Myers et al., 2011; Robles and Gomez, 2006; Woo et al., 2009). Both cell motility and growth cone pathfinding are triggered by guidance cues that activate cellular signaling mechanisms. Following exposure to growth and guidance cues, kinases, such as Src and FAK, are activated, thereby affecting adhesion formation (llić et al., 1995; Myers and Gomez, 2011; Robles and Gomez, 2006; Volberg et al., 2001). During adhesion formation, talin and paxillin are recruited first. Talin is a large adapter protein that can simultaneously bind both integrins and actin, thereby linking them. Paxillin is a scaffolding protein that directly interacts with integrins and recruits and facilitates interactions with other intracellular adhesion proteins (Lopez-Colome et al., 2017; Young et al., 2001). Following talin and paxillin recruitment, vinculin then binds to talin and strengthens the link of talin with both integrins and actin (Ciobanasu et al., 2014; del Rio et al., 2009). Meanwhile, the ribosomal binding protein, RACK1, which is regulated by Src and FAK, also localizes to adhesions (Adams et al., 2011; Duff and Long, 2017). RACK1 can act as a scaffold that promotes FAK phosphorylation, assisting in the recruitment of other proteins to adhesion sites (Kiely et al., 2009). Previous work has shown that when RACK1 interactions are suppressed, FAK signaling is dysregulated, leading to aberrant cell motility (Kiely et al., 2005).

Interestingly, previous sequencing studies using pathway analyses have shown that cellular adhesion is one of the most dysregulated biological processes in T21. Specifically, mRNAs encoding adhesion-related proteins, including paxillin, vinculin, and talin, have been shown to be upregulated in Down syndrome (Gonzales et al., 2018; Huo et al., 2018; Rastogi et al., 2024; Sobol et al., 2019; Susco et al., 2022). However, whether corresponding protein levels are also altered has not been examined, and changes in RNA abundance are often not reflected at the protein level (Srivastava et al., 2022; Vogel and Marcotte, 2012).

In this study, we used both fibroblasts and human induced pluripotent stem cell (hiPSC)- derived cortical neurons from individuals with Down syndrome to examine cellular morphology, migration, and adhesion proteins. In addition to their direct contribution to certain clinical features of Down syndrome, fibroblasts are a valuable model for examining cellular and molecular processes underlying axon growth and guidance during neuronal development, as these two cell types use similar motility mechanisms. When examining data from both models, we find that altered cellular morphology and motility are conserved features of Down syndrome. However, the direction of these changes, as well as changes in cellular adhesion proteins, is variable from individual to individual. These data show that dysregulated cellular morphology and motility may contribute to numerous phenotypes of Down syndrome, but the underlying mechanisms causing these phenotypes are heterogeneous.

## MATERIALS & METHODS

### Human Fibroblast Cell Culture

Three cell lines obtained from individuals with Down syndrome and age- and sex-matched control individuals were purchased from the Coriell Institute for Medical Research Biobank (GM05659 (Con.1); AG05397 (T21.1); AG07095 (Con.2); AG06922 (T21.2); GM05381 (Con.3); AG04823 (T21.3)). Cells were grown on 100-mm culture dishes (USA Scientific; CC7682-3394), maintained at 37°C in a humidified 5% CO2 incubator, and grown to 70-90% confluency in DMEM (VWR, 16750-074) supplemented with 10% Fetal Bovine Serum (FBS; Sigma F-2442) and 1% penicillin and streptomycin (pen/strep; Thermo Fisher 15140-122). Cells were counted using a hemocytometer, and 6,000 cells were plated onto a glass coverslip. Prior to plating, coverslips (Carolina; 633031) were submerged in nitric acid for 48 hours. Coverslips were then washed with deionized water four times for 30 minutes each before being baked for 3 hours at 120°C. After baking, coverslips were then stored in sterile petri dishes until further use. After sterilization, coverslips were coated with Matrigel (Corning, 47743-716) in serum-free DMEM (VWR, 16750-074) at a 1:100 dilution for 24 hours. After 24 hours, coverslips were washed with serum-free DMEM and replaced with DMEM supplemented with 10% Fetal Bovine Serum (FBS; Sigma, F-2442) and 1% penicillin and streptomycin (pen/strep; Thermo Fisher 15140-122) prior to plating. 24 hours after plating, fibroblasts were fixed with freshly made 4% paraformaldehyde (PFA) with 5mM MgCl_2_ (VWR, 100504-858). Cells were washed with 1X PBS (with 2mM MgCl2, pH 7.4) three times for five minutes each. Following fixation, cells were stored at 4°C up to one week before performing immunocytochemistry.

### Maintenance of human induced pluripotent stem cells (hiPSCs)

Human induced pluripotent stem cells (hiPSCs) from an individual with Down syndrome and an age- and sex-matched control were obtained from Dr. Alberto Costa (Case Western University; Cleveland, Ohio) (Lee et al., 2017). All cells were cultured on Nunc-coated tissue culture-treated dishes (VWR, 73520-906) coated with Matrigel (Corning, 354277), per manufacturer’s specifications. Media changes on hiPSCs were performed daily with fresh mTeSR1 media (Stem Cell Technologies, 85857). Every 6-7 days, hiPSCs were passaged using Gentle Cell Dissociation Reagent (Stem Cell Technologies, 100-0485). All wells were checked daily for spontaneous differentiation and scraped when spontaneous differentiation exceeded 10% of the dish.

### Generation of hiPSC-derived cortical neurons

Human iPSCs were differentiated using dual SMAD inhibition in monolayer culture as previously described (Agrawal et al., 2026; Shi et al., 2012; Volpato et al., 2018). This protocol uses Neuronal Maintenance Media (NMM). NMM is composed of a 1:1 ratio of DMEM/F-12 GlutaMAX (Gibco, 10565-018): Neurobasal (Gibco, 12348-017), 1x N2 supplement (Gibco, 17502001), 1x B27 supplement (Gibco, 17504001), 5 μg/mL insulin (Sigma, I9278), 1 mM L-glutamine (Gibco, 25030081), 500 μM sodium pyruvate (Sigma, S8636), 100 μM nonessential amino acids solution (Gibco, 11140050), 100 μM beta-mercaptoethanol (Gibco, M3148), 50 U per mL penicillin-streptomycin (Gibco, 15140-122). To set up for neuronal differentiation, hiPSCs from 2 wells, about 80-90% confluency each, were passaged into one well of a matrigel-coated 6-well plate to have a 100% confluency the following day (Day 0). The next day (Day 0), Neuronal Induction Media (NIM) was added, which contains NMM,1 μM Dorsomorphin (Tocris, 309310), and 10 μM SB431542 (Reprocell, 04-0010). Cells were maintained in NIM for 12 days with media changes every day to allow for the formation of the neuroepithelial sheet. On day 12, cells were passaged by gently breaking up the neuroepithelial sheet using a split ratio of 1:2 with 1mg/mL dispase (Sigma, D4693), and plating onto laminin-coated 6-well dishes. The following day (Day 13), media was changed to NMM supplemented with 20 ng/mL FGF2 (Gibco, PHG0024), with media changes every alternate day. After 4 days in NMM with 20 ng/mL FGF2 (Day 16), cells were maintained in NMM with media changes every alternate day and passaged at a 1:2 ratio following the growth of neuronal rosettes. At Day 25 after neuronal induction, cultures were passaged 1:1 into single cells using Accutase (Invitrogen, 00-4555-56) onto laminin-coated 6-well plates. Once wells were about 70-80% confluent, cells were then passaged at a 1:2 split ratio every 2-3 days using Accutase until about Day 35. At Day 35-40, cells were passaged using Accutase for final plating. 10,000 cells were plated onto each glass coverslip (washed as described above; Carolina Biological, 633031) pre-coated with 100 µg/mL poly-l-lysine (Sigma-Aldrich, P1274) and 10 µg/mL laminin (Life Technologies, 23017015). The next day, media was replaced with fresh NMM, and cells were cultured for an additional 2-3 days *in vitro* (DIV) before fixation. For studies examining adhesion proteins at basal conditions, hiPSC-derived neurons were fixed in freshly made 4% PFA in 1x PBS (with 4% sucrose, 5 mM MgCl2, pH 7.4). Cells were washed with 1X PBS (with 2mM MgCl2, pH 7.4) three times for five minutes each. Following fixation, cells were stored at 4°C up to one week before performing immunocytochemistry.

When stimulating hiPSC-derived neurons in bath application, neurons were allowed to grow for an additional 2-3 DIV in NMM after final plating. After 2-3 DIV, cells were starved by removing growth factors in the media (NMM-0; 1:1 ratio of DMEM/F-12 GlutaMAX: Neurobasal, 5 μg/mL insulin, 500 μM sodium pyruvate (Sigma, S8636), 100 μM beta-mercaptoethanol, 50 U per mL penicillin-streptomycin) for 3 hours (Kershner and Welshhans, 2017; Sasaki et al., 2010). After 3 hours, cells were stimulated with 100 ng/ml BDNF (PeproTech, 450-02) in NMM for 20 minutes before being fixed in 4% PFA in 1x PBS (with 4% sucrose, 5 mM MgCl2, pH 7.4). Cells were then washed with 1X PBS (with 2mM MgCl2, pH 7.4) three times for 5 minutes.

### Immunocytochemistry

Following the above fixation protocols, coverslips were washed with 1x TBS_50_ (50 mM Tris/HCl, 150 mM NaCl, pH 7.4) for 5 minutes. Cells were then permeabilized in 0.3% Triton x-100 (Sigma Aldrich, T8787) in 1x TBS_50_ for 5 minutes. Following permeabilization, cells were washed for 5 minutes in IF buffer (0.1% Triton x-100, 2% BSA, in 1x TBS_50_) before going into blocking buffer (0.1% Triton x-100, 2% BSA, 2% FBS, in 1x TBS_50_) for 1-2 hours. Coverslips were incubated with the following primary antibodies: rabbit anti-paxillin (1:500; Abcam, Ab32084), mouse anti-vinculin (1:500; Sigma, V9131), rabbit anti-talin (1:200; Abcam, Ab71333), and mouse anti-RACK1 (1:500; Santa Cruz, sc-17754) for 1 hour at room temperature or overnight at 4°C. Following primary incubation, coverslips were washed with IF buffer four times for 5 min. Coverslips were then incubated with secondary antibodies: donkey anti-rabbit Alexa 488 (1:1000; Life Technologies, A11034) and goat anti-mouse Alexa 568 (1:1000; Life Technologies, A11031) for 30 minutes. Following secondary incubation, coverslips were washed an additional 4 times with IF buffer for 5 minutes per wash. All coverslips were then washed with molecular biology-grade water and mounted onto glass slides (VWR, 48311- 951) using ProLong Gold anti-fade mounting media (Invitrogen, P36934). All slides were dried in the dark overnight prior to imaging.

### Transwell Membrane Assay

Fibroblasts were grown on 100-mm culture dishes (USA Scientific, CC7682-3394) until reaching 70-90% confluency. Then, cells were starved for 3 hours in serum-free media without fetal bovine serum. Following starvation, 10,000 cells were plated onto 8 µm pore size transwell inserts (Thincerts, VWR, 82050-038) in starvation media and allowed to settle for 10 minutes. DMEM with 10% FBS was added to the bottom of the well, and cells were allowed to migrate through the membrane for 24 hours. Using a cotton-tipped applicator, any non-migrated cells were wiped from the inner membrane, and migrated cells were fixed immediately in freshly made 4% PFA in Krebs Buffer (145 mM NaCl, 5 mM KCl, 1.2 mM CaCl2, 1.3 mM MgCl2, 1.2 mM NaH2PO4, 10 mM glucose, 20 mM HEPES, pH 7.4) for 17 minutes.

Immunocytochemistry was performed on 4% PFA-fixed fibroblasts. Inserts were first washed with 1X PBS 3 times, 5 minutes each. Next, inserts were washed in a permeabilization buffer made of 1X PBS, 0.3% Triton X-100 (Sigma Aldrich, T8787), 4% BSA (Sigma Aldrich, 10711454001), and 10% Goat Serum (VWR, 0219135680) for 15-20 minutes. The inserts were then washed 3 times for 5 minutes each with 0.5% BSA in 1X Krebs buffer as previously described (Dent and Meiri, 1992; Ghate et al., 2020). After washing, the inserts were submerged in a blocking buffer (3% BSA in 1X PBS) for 1 hour. Inserts were then incubated with acti-stain 488 phalloidin (7:1000; Cytoskeleton, PHDG1) and DAPI (1:1000; Sigma, A11031) in 1 mL blocking buffer (4% BSA, 10% Goat Serum in 1x PBS) overnight at 4°C. The next day, inserts were washed 3 times for 5 minutes each with 0.5% BSA in 1x Krebs buffer. Immediately thereafter, the hooks on the inserts were cut off using scissors, and the insert was placed in a 12-well glass bottom plate on a drop of molecular biology-grade water (VWR, 45001-044) for imaging.

### Image Acquisition and Analysis

All images were acquired using a Nikon Ti2-E microscope and Hamamatsu ORCA-Fusion camera. Fluorescence was visualized and captured, keeping all acquisition parameters consistent across all experimental groups. Fluorescence intensity was quantified using the image analysis software, Fiji.

The transwell membrane assay was imaged using a 10X objective. Five regions of interest (ROIs) were imaged per insert using a tile scan. All migrated cells were counted using the cell counter plugin in Fiji.

For all fibroblast morphology and immunofluorescence experiments, fluorescence images were captured on the 20X objective with a 1.5X multiplier for a final magnification of 30X. All growth cone images were taken using the 100X objective. Fluorescence intensity and area were quantified using the polygon tool to outline the entire perimeter of the fibroblast/growth cone in DIC. Fluorescence was quantified by measuring the mean grey value and subtracting the background intensity in Fiji. Perimeter was quantified using the segmented line tool in Fiji by outlining the entire perimeter of the cell. Length/Width ratio was measured using the straight-line tool in Fiji by measuring the length along the major axis of the cell (spanning the leading edge to the lagging edge) and dividing it by the width, which was measured along the minor axis at its widest point. Axon length was quantified using the segmented line tool by tracing the length of the longest neurite from the most proximal portion of the axon down to the tip of the growth cone.

All Puncta analysis was performed in Fiji by thresholding all images using the ‘RenyiEntropy’ algorithm to avoid individual bias. After thresholding, a mask of the puncta was created for each image. Under the ‘Analyze’ tab in FIJI, ‘Analyze Particles’ was used to create ROIs of individual puncta in the ‘ROI Manager’. In the ‘ROI Manager’, selecting individual ROIs created an outline around the puncta in the original image. Next, ‘Measure’ in ‘ROI Manager’ was used to quantify the area and mean fluorescence intensity of each selected puncta.

Obvious clusters of puncta that the algorithm failed to identify as distinct puncta were excluded from analyses. Puncta per area was then calculated by dividing the total puncta in a single growth cone by the area of that growth cone. The area of the growth cone was calculated by outlining growth cones using the polygon tool in Fiji.

Pearson’s Correlation Coefficient was quantified in growth cones of hiPSC-derived neurons using Just another Co-localization Plugin (JaCoP) in Fiji software (Bolte and Cordelieres, 2006). Pearson’s Correlation coefficient measures the linear correlation between two variables, with values ranging from -1 to 1, and 1 signifying perfect correlation (Dunn et al., 2011). ROIs of growth cones were marked using the box tool, before being duplicated into a new window. All image stacks were split into individual channel images and selected in the JaCoP window for colocalization analyses.

### Statistical Analysis

All statistical analyses were conducted in GraphPad Prism software with a significance value set at p ≤ 0.05. All data were first tested for normality, and the appropriate statistical test was then applied based on that result. Specific statistical tests for each experiment are given in the figure legend. Error bars in the figures represent the standard error of the mean of three independent passages or differentiation sets unless otherwise stated.

### Study Approval

All work with iPSCs was reviewed by the Institutional Review Board at the University of South Carolina and designated as exempt.

## RESULTS

### Morphology is dysregulated in Down syndrome fibroblasts

We first used three distinct pairs of fibroblasts obtained from individuals with Down syndrome and matched controls to investigate whether trisomy 21 alters cellular morphology. Pair 1 (Con.1 and T21.1) consisted of skin fibroblasts from male individuals at 1 year of age; Pair 2 (Con.2 and T21.2) consisted of skin fibroblasts from male individuals at 2 years of age; and Pair 3 (Con.3 and T21.3) consisted of skin fibroblasts from male individuals at 5 years of age. First, we quantified the area, perimeter, and length-to-width ratio of the cells (**Figure 1**). Pairs 1 and 3 showed a significant increase in the area of the T21 fibroblasts compared with controls (**Figure 1A-B & E; Con.1 vs. T21.1 and Con.3 vs. T21.3**). However, T21.2 showed a significant decrease in area compared with control (**Figure 1A-B & E.2 vs. T21.2**). Perimeter was increased in T21.3 fibroblasts compared with control, but Pair 1 was not significantly different (**Figure 1C & F**). Additionally, the perimeter of T21.2 was decreased, following the same trend quantified in area (**Figure 1B-C, E-F**). Lastly, we measured the length-to-width ratio (Length/Width) to compare the size proportion of these cells. There was a significant decrease in the length-to-width ratio in T21.3 fibroblasts compared with control (**Figure 1D & G**).

**Figure 1.**
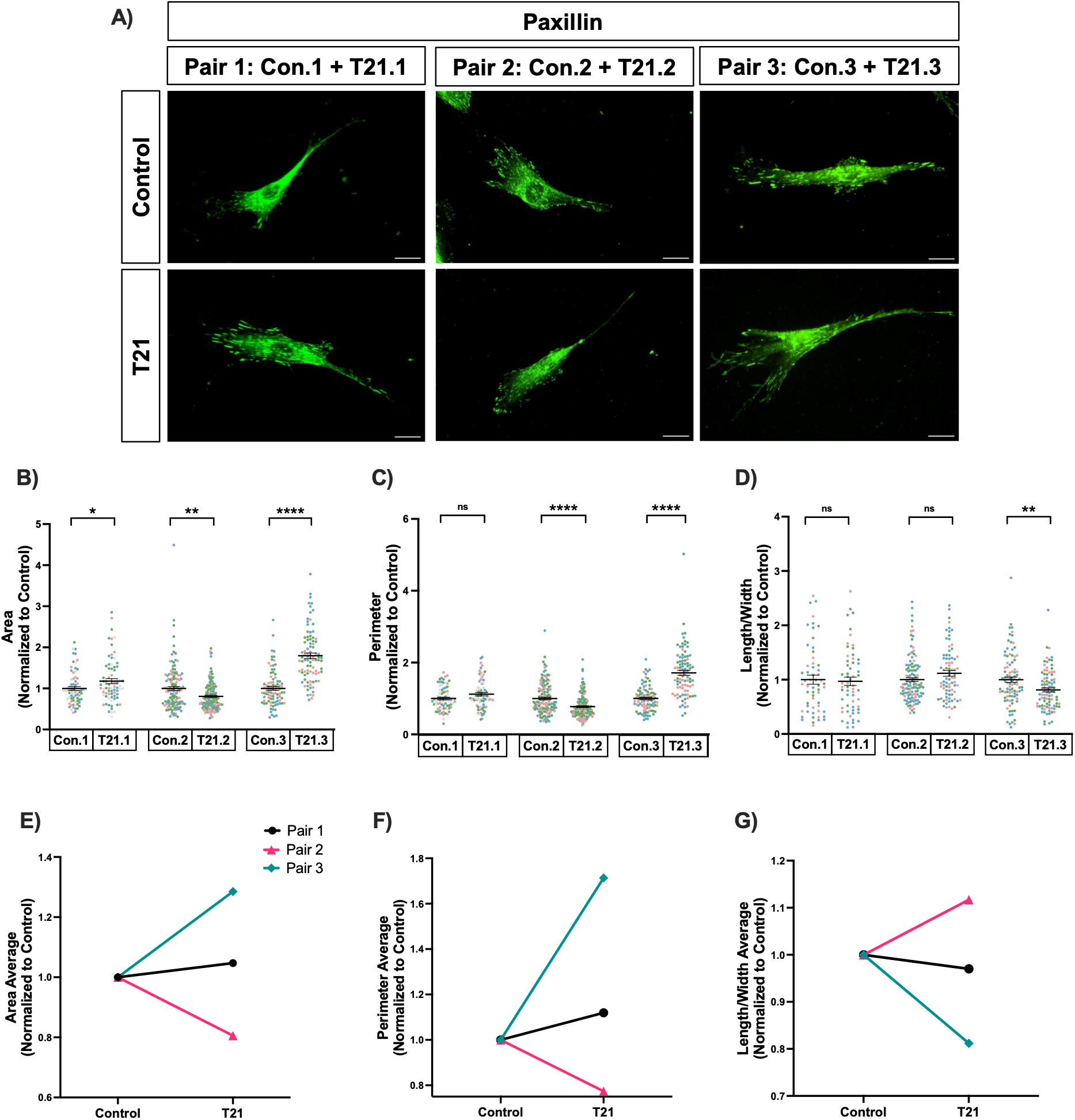
Morphology is dysregulated in T21 fibroblasts. **(A)** Representative images of Down syndrome (T21.1, T21.2, T21.3) and control (Con.1, Con.2, Con.3) fibroblasts stained for the focal adhesion protein, paxillin. Scale Bars, 25 µm. **(B&E)** Fibroblast area was quantified and shown as individual data points with mean ± SEM (B) and overall averages for each pair (E). Each experiment was repeated 3 independent times. Each replicate is signified by a separate color. Con.1: n=73, T21.1: n=75; *p=0.0288, Mann-Whitney. Con.2: n=133, T21.2: n=147; **p=0.0015, Mann-Whitney. Con.3: n=90, T21.3: n=90; ****p<0.0001, Mann-Whitney. **(C&F)** Fibroblast perimeter was quantified and shown as individual data points with mean ± SEM (C) and overall averages for each pair (F). Each experiment was repeated 3 independent times. Each replicate is signified by a separate color. Con.1: n=75, T21.1: n=76; p=0.0678, Mann-Whitney. Con.2: n=133, T21.2: n=147; ****p<0.0001, Mann-Whitney. Con.3: n=95, T21.3: n=92; ****p<0.0001, Mann-Whitney. **(D&G)** The length-to-width ratio of fibroblasts was quantified and shown as individual data points with mean ± SEM (D) and overall averages for each pair (G). Each experiment was repeated 3 independent times. Each replicate is signified by a separate color. Con.1: n=57, T21.1: n=59; p=0.9124, Mann-Whitney. Con.2: n=124, T21.2: n=80; p=0.0736, Mann-Whitney. Con.3: n=95, T21.3: n=92; **p=0.0033, Mann-Whitney.

However, there was no significant difference in the length-to-width ratio for either Pair 1 or 2 (**Figure 1D & G)**. Together, these data suggest that although all T21 fibroblasts showed significant changes in cellular morphology, the direction of this change varies between individuals.

### Motility is often decreased in T21 fibroblasts

Given the changes in cell size, we reasoned that migration may also be affected. Thus, a transwell membrane assay was used to quantify cell motility in all three fibroblast pairs. Cells were starved for 3 hours in serum-free DMEM to induce a more quiescent state. Following the starvation period, 10,000 cells were plated on top of a transwell membrane in serum-free DMEM. At the bottom of the chamber, DMEM containing serum (10% FBS) was added, and cells were allowed to migrate through the membrane, toward serum-containing media, for 24 hours (Justus et al., 2014; Omar Zaki et al., 2019). In line with the increase in cellular area (**Figures 1B & E**), fewer T21.1 and T21.3 cells migrated through the membrane, compared with control. However, Pair 2 showed no significant difference in motility (**Figure 2A-B**). Together, these data suggest that cellular motility is often altered in T21.

**Figure 2.**
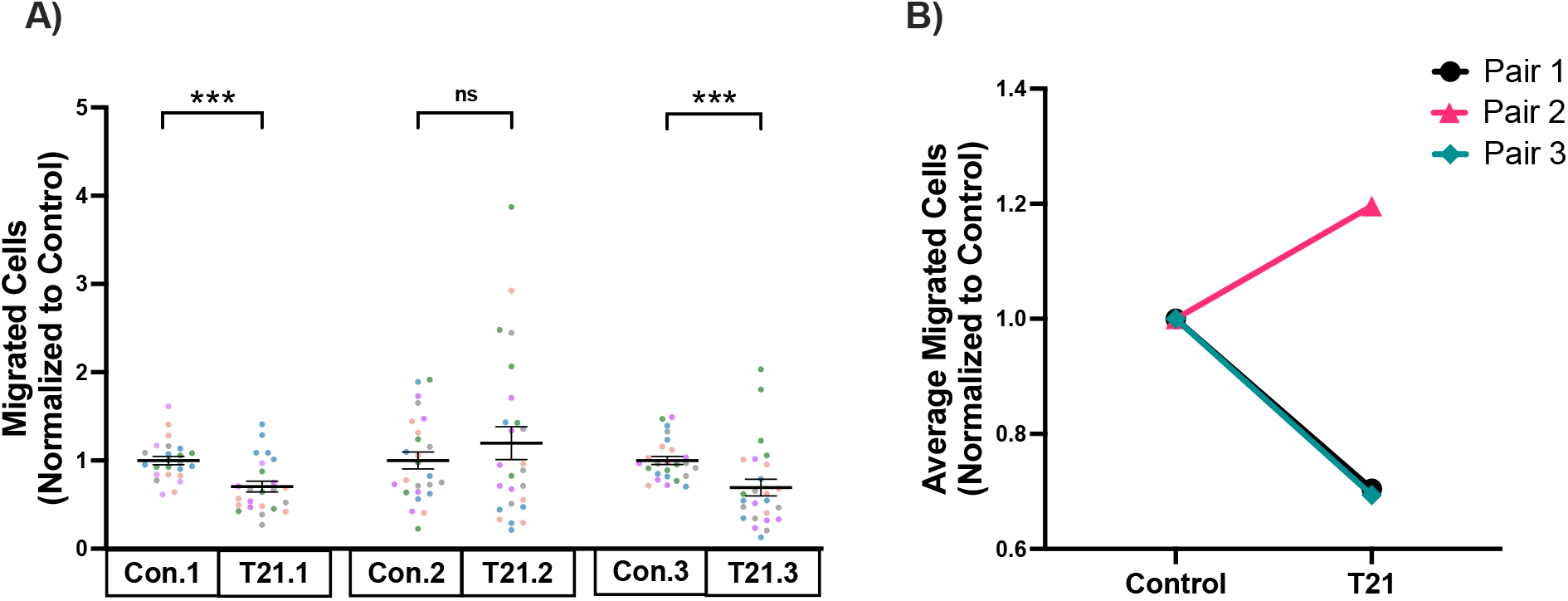
Motility is decreased in T21 fibroblasts. **(A&B)** Trisomy 21 (T21.1, T21.2, T21.3) and control (Con.1, Con.2, Con.3) fibroblasts were starved for 3 hours in serum-free DMEM, plated on the top of transwell membranes in serum-free DMEM, and allowed to migrate through the membrane toward FBS containing media for 24 hours. The number of cells that migrated through the membrane after 24 hours was quantified and is shown as individual data points with mean ± SEM (A) and overall averages for each pair (B). Each experiment was repeated 5 independent times, with each replicate shown in a separate color. Con.1: n=25, T21.1: n=25; ***p=0.0004, Unpaired t-test. Con.2: n=25, T21.2: n=25; p=0.3543, Unpaired t-test. Con.3: n=25, T21.3: n=25; ***p=0.0005, Mann-Whitney.

### Adhesion proteins are dysregulated in T21 fibroblasts

The formation and turnover of focal adhesion sites regulate actin cytoskeletal dynamics and are necessary for cell morphology and motility (Ciobanasu et al., 2012). Thus, quantitative immunofluorescence was used to label four core adhesion proteins (paxillin, vinculin, talin, and RACK1) and quantify mean fluorescence intensity in T21 and control fibroblasts (**Figure 3A**).

**Figure 3.**
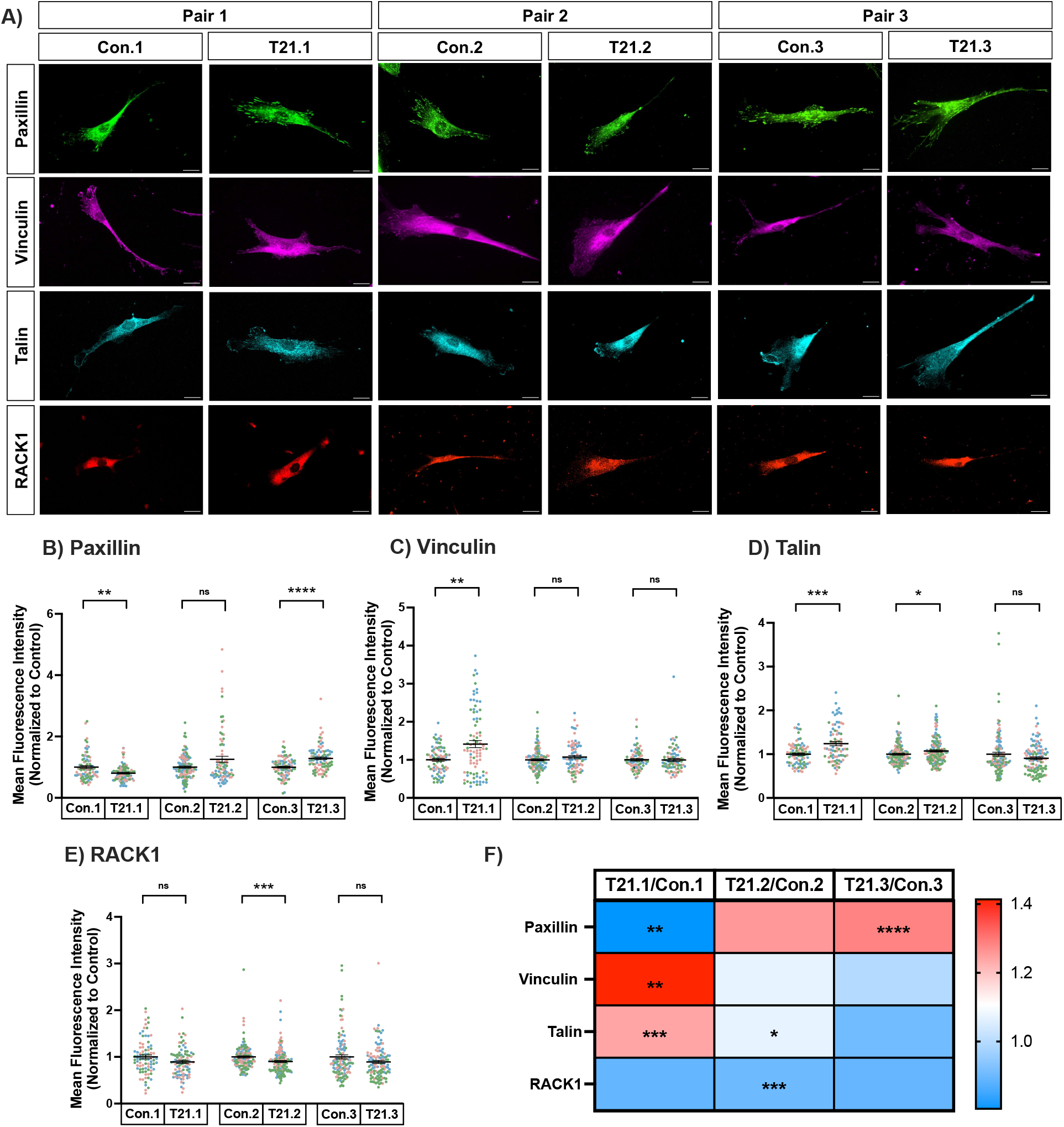
Adhesion proteins are dysregulated in T21 fibroblasts. **(A)** Representative images of trisomy 21 (T21.1, T21.2, T21.3) and control (Con.1, Con.2, Con.3) fibroblasts stained for the focal adhesion proteins paxillin, vinculin, talin, and RACK1. Note that the RACK1 cells appear smaller due to more intense RACK1 immunostaining in the cell center and weaker staining in the periphery as compared to the other adhesion proteins examined. Scale Bars, 25 µm. **(B-E)** Quantitative immunofluorescence (QIF) for paxillin (B), vinculin (C), and talin (D), and RACK1 (E) was performed and shown as individual data points with mean ± SEM. (B) Con.1: n=84, T21.1: n=88; **p=0.0014, Mann-Whitney. Con.2: n=124, T21.2: n=80; p=0.6688, Mann-Whitney. Con.3: n=90, T21.3: n=90; ****p<0.0001, Mann- Whitney. (C) Con.1: n=85, T21.1: n=85; **p=0.0086, Mann-Whitney. Con.2: n=124, T21.2: n=80; p=0.3218, Mann-Whitney. Con.3: n=78, T21.3: n=80; p=0.5312, Mann-Whitney. (D) Con.1: n=87, T21.1: n=79; ***p=0.0002, Mann-Whitney. Con.2: n=133, T21.2: n=147; *p=0.0308, Mann-Whitney. Con.3: n=121, T21.3: n=117; p=0.3293, Mann-Whitney. (E) Con.1: n=84, T21.1: n=85; p=0.0518, Mann-Whitney. Con.2: n=133, T21.2: n=147; ***p=0.0001, Mann- Whitney. Con.3: n=121, T21.3: n=117; p=0.1584, Mann-Whitney. **(F)** Heat map comparison summarizes the mean fluorescence intensity of paxillin, vinculin, talin, and RACK1 in the 3 pairs of T21 fibroblasts compared to control, respectively. All experiments were repeated 3 independent times. Each replicate is shown in a separate color.

Paxillin was significantly reduced in T21.1 cells and significantly increased in T21.3 cells compared with control. Pair 2 did not show a significant difference in paxillin, but the values were more variable in T21.2 (SEM: 1.257 ± 0.0985) and showed a trend toward an increase (**Figure 3B & F**). Vinculin was significantly increased in Pair 1 T21 cells compared with control, but no changes were detected in Pair 2 or 3 (**Figure 3C & F**). Talin was significantly increased in both T21.1 and T21.2 compared with their respective controls, whereas Pair 3 showed no difference (**Figure 3D & F**). Lastly, RACK1 was significantly decreased in Pair 2, but there was no significant difference between T21 and control cells for Pair 1 and Pair 3 (**Figure 3E-F**). In summary, cell morphology, cell motility, and adhesion proteins are altered in T21 fibroblasts, but the proteins that are altered and the extent of their protein expression changes vary among individuals (**Supplemental Table 1**).

### Growth cone area and the longest neurite length are decreased in T21 hiPSC-derived cortical neurons

To determine whether the cellular morphology changes observed in fibroblasts also occur in neurons, we examined hiPSC-derived cortical neurons (**Figure 4**). hiPSCs from an individual with Down syndrome and an age- and sex-matched control were cultured and then differentiated into cortical neurons over 35-40 DIV before being plated on coverslips (Agrawal et al., 2026; Lee et al., 2017; Shi et al., 2012; Volpato et al., 2018). Two to three days after final plating, when one neurite had elongated extensively to form the axon (Stage 3 of neuritogenesis) (Govek et al., 2005), the cells were fixed, and growth cone area and the length of the longest neurite were quantified. Both growth cone area and the length of the longest neurite were significantly decreased in T21 hiPSC-derived cortical neurons (**Figure 4A-C**). These data are consistent with a previous study from our lab, which demonstrated that soma area, neurite branching, number of neurites, and attractive axon guidance are decreased in T21 hiPSC-derived cortical neurons (Agrawal et al., 2026).

**Figure 4.**
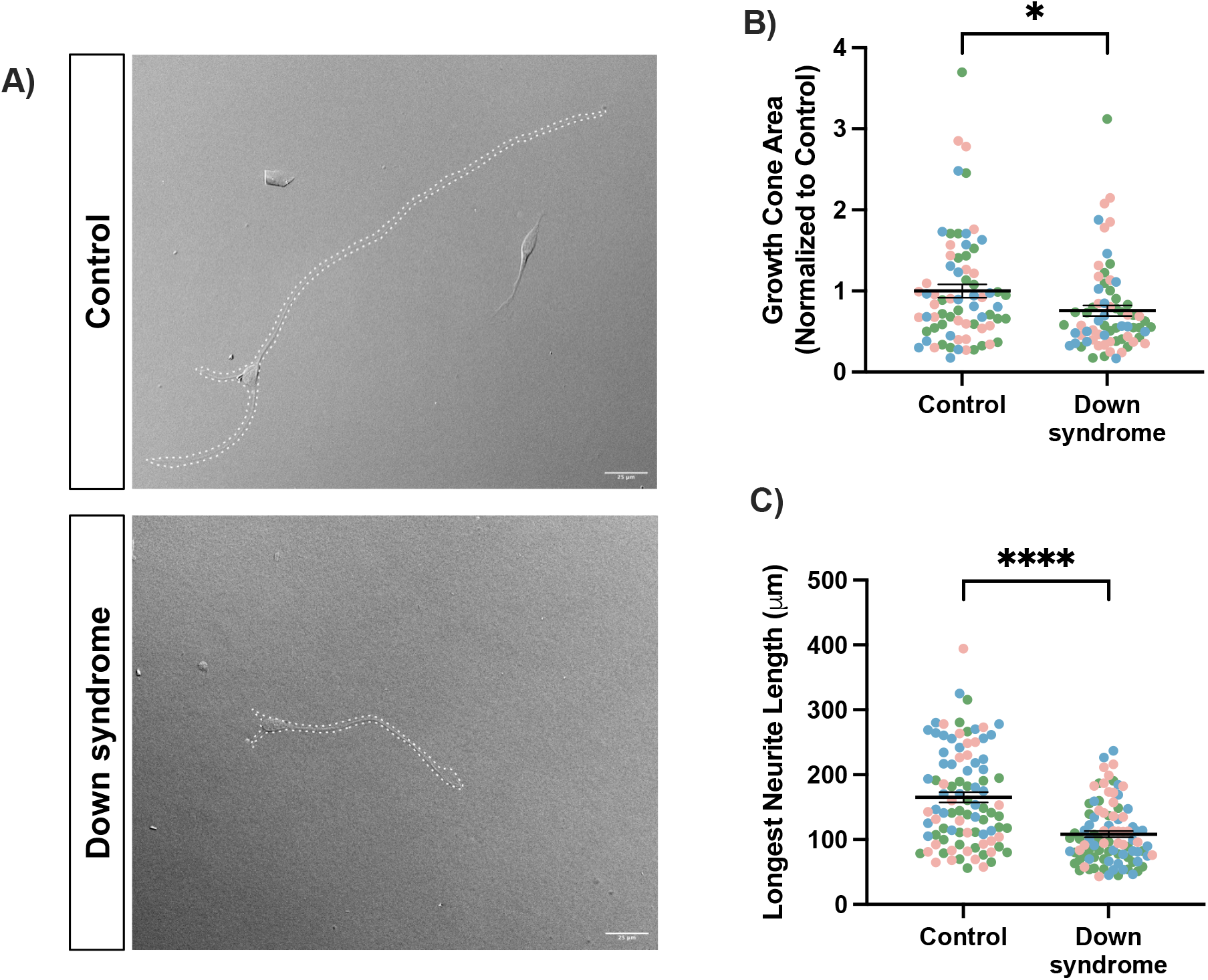
Growth cone area and the longest neurite length are decreased in Down syndrome hiPSC-derived cortical neurons. **(A)** Representative images of neurite length in control and Down syndrome hiPSC-derived derived cortical neurons. The dotted line outlines the entire cell, but only the growth cone area (B) and the length of the longest neurite (C) were quantified. Scale bars 25μm. **(B)** Growth cone area of control and Down syndrome hiPSC- derived cortical neurons was quantified and shown as individual data points with mean ± SEM. *p=0.0134, Mann-Whitney. Control n=71 growth cones; Down syndrome n=69 growth cones. **(C)** The length of the longest neurite in control and Down syndrome hiPSC-derived cortical neurons was quantified and shown as shown as individual data points with mean ± SEM. ****p<0.0001, Mann-Whitney. Control n=93 neurons; Down syndrome n=94 neurons. “n” is the number analyzed over 3 independent differentiations. Each differentiation set is signified by a separate color.

### Adhesion proteins talin and RACK1 are dysregulated in growth cones of T21 hiPSC- derived cortical neurons under basal conditions

Because we have previously shown that multiple morphological parameters are decreased in T21, including axon length, growth cone area, axon branching, and attractive axon guidance (Figure 4 and Agrawal et al., 2026), we examined whether adhesion proteins that regulate these processes may also be altered. After final plating, we allowed cells to grow for an additional 2 DIV before performing quantitative immunocytochemistry to fluorescently label adhesion proteins under basal conditions (**Figure 5A**). First, the mean fluorescence intensity of each protein in growth cones and the number of puncta per growth cone area (Puncta/Area) were quantified. Because previous work has suggested that larger adhesion sites correlate with stronger or less dynamic adhesions in non-neuronal cells, the area and mean fluorescence intensity of each individual puncta within the growth cones were also quantified (Schiller and Fassler, 2013; Schumacher et al., 2022). Paxillin did not differ significantly in T21 growth cones compared to control across these four parameters (**Figure 5B.i-B.iv**). Similarly, the mean fluorescence intensity, puncta per area, and puncta area of vinculin showed no significant differences between T21 and control growth cones (**Figure 5C.i-C.iii**). However, the fluorescence intensity of vinculin puncta was decreased in T21 growth cones compared to control (**Figure 5C.iv**). For talin and RACK1, the mean fluorescence intensity in T21 growth cones was increased, with no significant differences in puncta per area, puncta area, or the fluorescence intensity of individual puncta (**Figure 5D.i-D.iv** & **5E.iii-E.iv**). Thus, talin and RACK1 are increased in T21 growth cones, but this does not affect the number or size of the adhesions. Furthermore, vinculin localization to adhesions is slightly, but significantly, decreased. Taken together, these results show that there are some changes in adhesion proteins in growth cones of T21 hiPSC-derived neurons under basal conditions, which could affect cellular morphology and motility.

**Figure 5.**
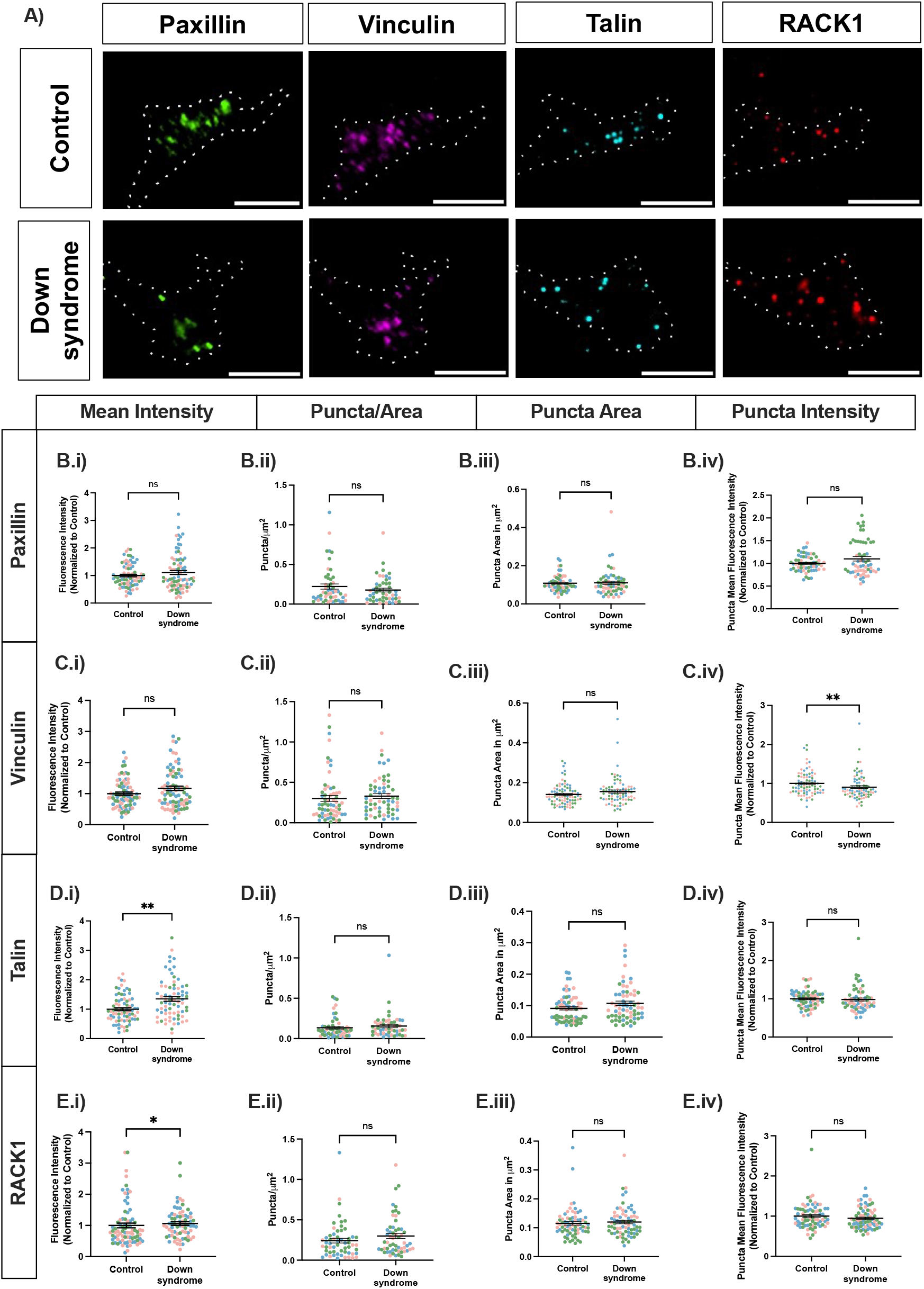
Vinculin, talin, and RACK1 are dysregulated in growth cones of Down syndrome hiPSC-derived neurons at basal conditions. **(A)** Representative images of growth cones of control and Down syndrome hiPSC-derived neurons stained for members of the adhesion complex: paxillin, vinculin, talin, and RACK1. Scale Bars, 5 µm. **(B-E)** Quantitative immunofluorescence for paxillin (B.i - B.iv), vinculin (C.i – C.iv), talin (D.i – D.iv), and RACK1 (E.i – E.iv) was performed, and shown as individual data points with mean ± SEM. (B.i, C.i, D.i, E.i) Mean fluorescence intensity within growth cones was quantified. (B.ii, C.ii, D.ii, E.ii) The number of puncta per area within growth cones was quantified. (B.iii, C.iii, D.iii, E.iii) The area of individual puncta in growth cones was quantified. (B.iv, C.iv, D.iv, E.iv) The intensity of individual puncta in growth cones was quantified. **(B.i)** p=0.9644, Mann-Whitney. Control n=80 growth cones; Down syndrome n=83 growth cones. **(B.ii)** p=0.518, Mann-Whitney. Control n=52 growth cones; Down syndrome n=49 growth cones. **(B.iii)** p=0.6073, Mann-Whitney. Control n=313 puncta from 60 growth cones; Down syndrome n=228 puncta from 60 growth cones. **(B.iv)** p=0.8651, Mann-Whitney. Control n=313 puncta from 60 growth cones; Down syndrome n=227 puncta from 60 growth cones. **(C.i)** p=0.2080, Mann-Whitney. Control n=84 growth cones; Down syndrome n=85 growth cones. **(C.ii)** p=0.1161, Mann-Whitney. Control n=64 growth cones; Down syndrome n=60 growth cones. **(C.iii)** p=0.0954, Mann-Whitney. Control n=554 puncta in 83 growth cones; Down syndrome n=650 puncta in 79 growth cones. **(C.iv)** **p=0.0081, Mann-Whitney. Control n=554 puncta from 83 growth cones; Down syndrome n=650 puncta from 79 growth cones. **(D.i)** **p=0.0012, Mann-Whitney. Control n=82; Down syndrome n=75. **(D.ii)** p=0.1708, Mann-Whitney. Control n=54 growth cones; Down syndrome n=50 growth cones. **(D.iii)** p=0.1366, Mann-Whitney. Control n=192 puncta from 65 growth cones; Down syndrome n=191 puncta from 64 growth cones. **(D.iv)** p=0.1489, Mann-Whitney. Control n=192 puncta from 65 growth cones; Down syndrome n=190 puncta from 64 growth cones. **(E.i)** *p=0.0341, Mann-Whitney. Control n=80; Down syndrome n=72. **(E.ii)** p=0.1521, Mann-Whitney. Control n=57 growth cones; Down syndrome n=57 growth cones. **(E.iii)** p=0.6169, Mann-Whitney. Control n=382 puncta from 71 growth cones; Down syndrome n=382 puncta from 72 growth cones. **(E.iv)** p=0.2400, Mann-Whitney. Control n=382 puncta from 71 growth cones; Down syndrome n=382 puncta from 72 growth cones. “n” is the number analyzed over 3 independent differentiations. Each differentiation set is signified by a separate color.

### RACK1 is dysregulated in growth cones of T21 hiPSC-derived cortical neurons under BDNF-stimulated conditions

In mouse cortical neurons, treatment with attractive guidance cues, such as brain-derived neurotrophic factor (BDNF), increases growth cone area and point contact number (Kershner and Welshhans, 2017). To investigate whether adhesion assembly may be affected in Down syndrome, we next examined adhesions following BDNF treatment. Control and T21 hiPSC- derived cortical neurons were cultured for 2 DIV after the final plating, then starved for three hours. Following starvation, neurons were treated with either a vehicle control or 100 ng/mL BDNF via bath application for 20 minutes. Neurons were stained for adhesion proteins using quantitative immunocytochemistry, and within growth cones, the following parameters were quantified for each adhesion protein: mean fluorescence intensity, puncta per area (number of puncta divided by growth cone area), area of each puncta, and the mean fluorescence intensity of each puncta (**Figure 6A**).

**Figure 6.**
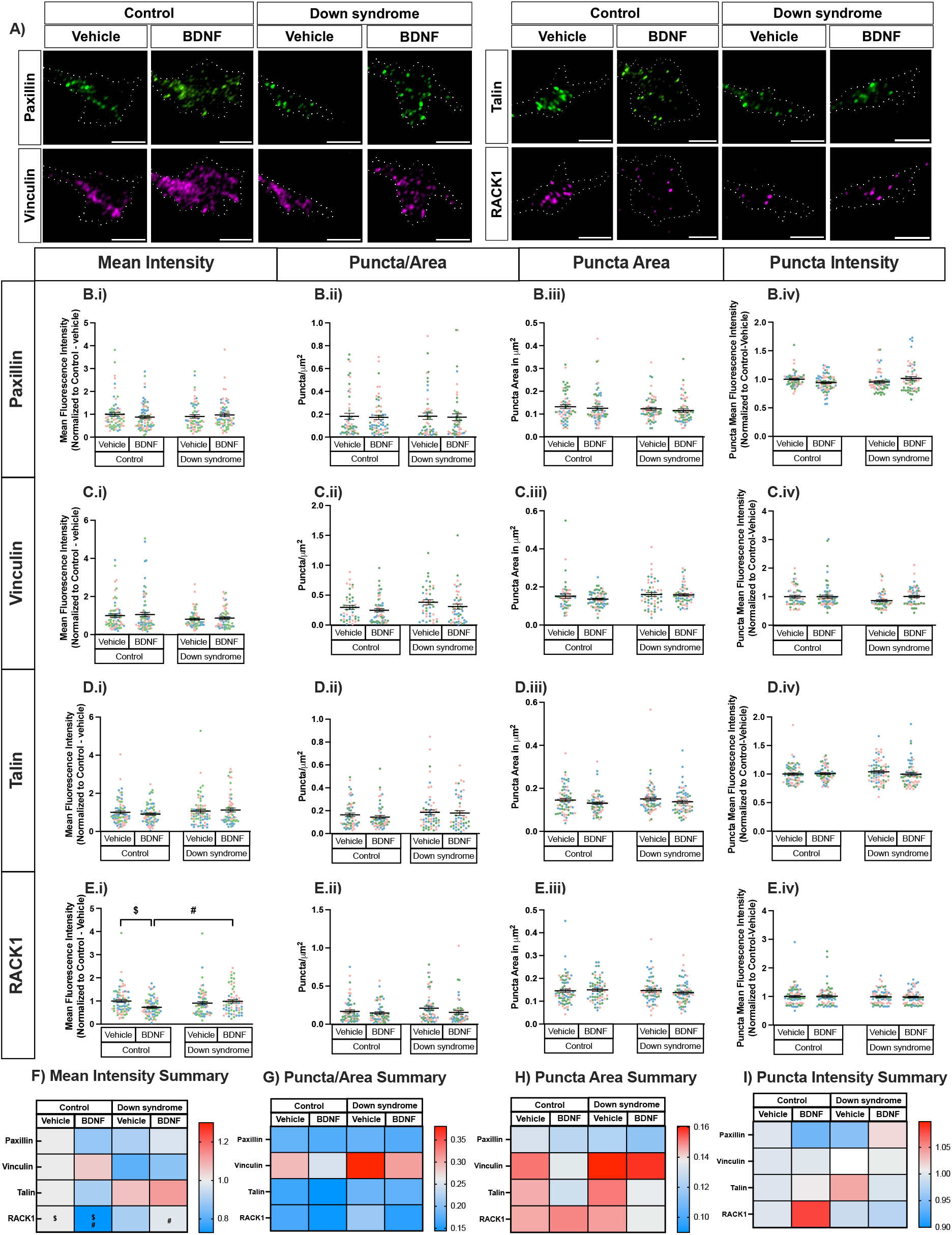
RACK1 is dysregulated in growth cones of Down syndrome hiPSC-derived cortical neurons under BDNF-stimulated conditions. **(A)** Representative images of growth cones on control and Down syndrome hiPSC-derived cortical neurons that were stimulated with 100ng/ml BDNF or vehicle, and stained for adhesion proteins (paxillin, vinculin, talin, and RACK1) using quantitative immunofluorescence. Scale bars, 5μm. **(B-E)** Quantitative immunofluorescence for paxillin (B.i - B.iv), vinculin (C.i – C.iv), talin (D.i – D.iv), and RACK1 (E.i – E.iv) was performed, and shown as individual data points with mean ± SEM. (B.i, C.i, D.i, E.i) Mean fluorescence intensity within growth cones was quantified. (B.ii, C.ii, D.ii, E.ii) The number of puncta per area within growth cones was quantified. (B.iii, C.iii, D.iii, E.iii) The area of individual puncta in growth cones was quantified. (B.iv, C.iv, D.iv, E.iv) The intensity of individual puncta in growth cones was quantified. **(B.i)** Two-way ANOVA: Genotype effect, p=0.9834; Treatment effect, p=0.6983; Interaction effect, p=0.2118. Control + Vehicle n=70; Control/BDNF n=77; Down syndrome + Vehicle n=60; Down syndrome/BDNF n=65. **(B.ii)** Two- way ANOVA: Genotype effect, p=0.9610; Treatment effect, p=0.7313; Interaction effect, p=0.9904. Control + Vehicle n=52; Control + BDNF n=61; Down syndrome + Vehicle n=50; Down syndrome + BDNF n=50. **(B.iii)** Two-way ANOVA: Genotype effect, p=0.2186; Treatment effect, p=0.3606; Interaction effect, p=0.9667. Control + Vehicle n=224 puncta from 58 growth cones; Control + BDNF n=323 puncta from 72 growth cones; Down syndrome + Vehicle n=239 from 61 growth cones; Down syndrome + BDNF n=251 from 58 growth cones. **(B.iv)** Two-way ANOVA: Genotype effect, p=0.5171; Treatment effect, p=0.9344; Interaction effect, **p=0.0068. Control + Vehicle n=224 puncta from 58 growth cones; Control + BDNF n=323 puncta from 72 growth cones; Down syndrome + Vehicle n=239 from 61 growth cones; Down syndrome + BDNF n=250 from 58 growth cones. **(C.i)** Two-way ANOVA: Genotype effect, *p=0.0352; Treatment effect, p=0.5300; Interaction effect, p=0.9715. Control + Vehicle n=70; Control/BDNF n=78; Down syndrome + Vehicle n=62; Down syndrome/BDNF n=65. **(C.ii)** Two-way ANOVA: Genotype effect, *p=0.0468; Treatment effect, p=0.1066; Interaction effect, p=0.7361. Tukey Post-Hoc: Control + BDNF vs. Down syndrome + Vehicle, *p=0.0474. Control + Vehicle n=43 growth cones; Control + BDNF n=58 growth cones; Down syndrome + Vehicle n=41 growth cones; Down syndrome + BDNF n=46 growth cones. **(C.iii)** Two-way ANOVA: Genotype effect, p=0.0588; Treatment effect, p=0.2849; Interaction effect, p=0.4229. Control + Vehicle n=339 puncta from 48 growth cones; Control + BDNF n=533 puncta from 68 growth cones; Down syndrome + Vehicle n=375 from 53 growth cones; Down syndrome + BDNF n=533 from 55 growth cones. **(C.iv)** Two-way ANOVA: Genotype effect, p=0.1836; Treatment effect, p=0.1414; Interaction effect, p=0.1433. Control + Vehicle n=341 puncta from 48 growth cones; Control + BDNF n=529 puncta from 68 growth cones; Down syndrome + Vehicle n=379 puncta from 53 growth cones; Down syndrome + BDNF n=410 puncta form 55 growth cones. **(D.i)** Two-way ANOVA: Genotype effect, p=0.0921; Treatment effect, p=0.8320; Interaction effect, p=0.3711. Control + Vehicle n=74; Control + BDNF n=71; Down syndrome + Vehicle n=67; Down syndrome + BDNF n=60. **(D.ii)** Two-way ANOVA: Genotype effect, p=0.1211; Treatment effect, p=0.4888; Interaction effect, p=0.7094. Control + Vehicle n=55; Control + BDNF n=54; Down syndrome + Vehicle n=61; Down syndrome + BDNF n=49. **(D.iii)** Two-way ANOVA: Genotype effect, p=0.4116; Treatment effect, p=0.0559; Interaction effect, p=0.9393. Control + Vehicle n=253 puncta from 68 growth cones; Control + BDNF n=252 puncta from 66 growth cones; Down syndrome + Vehicle n=266 puncta from 71 growth cones; Down syndrome + BDNF n=221 puncta from 63 growth cones. **(D.iv)** Two-way ANOVA: Genotype effect, p=0.5102; Treatment effect, p=0.4332; Interaction effect, p=0.2594. Control + Vehicle n=253 puncta from 68 growth cones; Control + BDNF n=252 puncta from 66 growth cones; Down syndrome + Vehicle n=265 puncta from 71 growth cones; Down syndrome + BDNF n=221 from 63 growth cones. **(E.i)** Two-way ANOVA: Genotype effect, p=0.2054; Treatment effect, p=0.1315; Interaction effect, **p=0.0067. Tukey Post-Hoc: Control + Vehicle vs. Control + BDNF, ^$^p=0.0113; Control + BDNF vs. Down syndrome + BDNF, ^#^p=0.0302. Control + Vehicle n=74 growth cones; Control + BDNF n=71 growth cones; Down syndrome + Vehicle n=68 growth cones; Down syndrome + BDNF n=60 growth cones. **(E.ii)** Two-way ANOVA: Genotype effect, p=0.2405; Treatment effect, p=0.0802; Interaction effect, p=0.4460. Control + Vehicle n=59 growth cones; Control + BDNF n=54 growth cones; Down syndrome + Vehicle n=57 growth cones; Down syndrome + BDNF n=54 growth cones. **(E.iii)** Two-way ANOVA: Genotype effect, p=0.4309; Treatment effect, p=0.6843; Interaction effect, p=0.3336. Control + Vehicle n=291 puncta from 73 growth cones; Control + BDNF n=224 puncta from 63 growth cones; Down syndrome + Vehicle n=300 puncta from 67 growth cones; Down syndrome + BDNF n=214 puncta from 65 growth cones. **(E.iv)** Two-way ANOVA: Genotype effect, p=0.5637; Treatment effect, p=0.9715; Interaction effect, p=0.8059. Control + Vehicle n=291 puncta from 73 growth cones; Control + BDNF n=224 puncta from 63 growth cones; Down syndrome + Vehicle n=300 puncta from 67 growth cones; Down syndrome + BDNF n=214 puncta from 65 growth cones. “n” is the number analyzed over 3 independent differentiations. Each differentiation set is signified by a separate color. **(F-I)** Heatmap summary of means of pairwise compairsons for fluorescence intensity (F), puncta/area (G), puncta area (H), and puncta fluorescence intensity (I) of paxillin, vinculin, talin, and RACK1.

For paxillin, there was no significant difference in mean fluorescence intensity, puncta number, puncta area, or puncta intensity per growth cone in T21 hiPSC-derived neurons compared to control (**Figure 6B.i-iv**). Vinculin was also not significantly different in any of these four parameters examined (**Figure 6C.i-iv**). However, the mean fluorescence intensity of vinculin was significantly decreased in T21 growth cones as compared to control (main effect of genotype; note that only post hoc analyses are shown on the graphs; **Figure 6C.i)**. Conversely, the number of vinculin puncta per area was significantly increased in T21 hiPSC-derived neurons (main effect of genotype; **Figure 6C.ii)**. Talin did not show any notable changes in T21 hiPSC-derived cortical neurons compared with control neurons (**Figure 6D.i-iv**). RACK1 was significantly decreased in control neurons following BDNF stimulation, but this decrease did not occur in T21 neurons (**Figure 6E.i**). The number of RACK1 puncta per area, puncta area, and puncta intensity were not significantly different when comparing T21 and control neurons (**Figure 6E.ii-iv**). Heat maps were used to summarize protein expression comparisons (**Figure 6F-I**). In summary, the mean intensity of RACK1 is altered in T21. However, changes in puncta number, area, and intensity of all of the adhesion proteins investigated were not significantly altered under vehicle or BDNF-stimulated conditions. However, it is important to note that, based on previous work using mouse models (Kershner and Welshhans, 2017), we expected a significant increase in all of the adhesion proteins in control hiPSC-derived neurons stimulated with BDNF, as compared to control neurons treated with vehicle; however, this did not occur and thus suggests differences in the response of developing mouse and human neurons to BDNF.

### The colocalization of adhesion proteins is not altered in T21 hiPSC-derived cortical neurons

Although we did not see widespread changes in individual adhesion proteins, adhesion composition is heterogeneous. Further, the protein composition of adhesions can affect how they function (Arregui et al., 1994; Renaudin et al., 1999; Wehrle-Haller, 2012; Woo and Gomez, 2006). To better understand the characteristics of these adhesion sites in Down syndrome following cortical neuron differentiation, we starved and stimulated both control and T21 hiPSC-derived cortical neurons with either vehicle or 100ng/ml BDNF by bath application for 20 minutes. Following fixation, neurons were stained for adhesion proteins, and a Pearson’s colocalization analysis was performed to measure colocalization. There were no significant differences in the colocalization of paxillin and vinculin between control and T21 neurons, either under basal or BDNF-stimulated conditions (**Figure 7A-B**). Colocalization analysis was also performed for talin and RACK1, and there were no significant differences between control and T21 neurons, either under basal or BDNF-stimulated conditions (**Figure 7C-D)**. Given the lack of statistically significant findings in Figures 6 and 7, we did not pursue additional colocalization combinations. Taken together, cellular morphology is altered in T21; however, there are only minor changes in adhesion protein expression, and point contact composition is not altered, in growth cones of T21 hiPSC-derived neurons (**Supplemental Table 2**).

**Figure 7.**
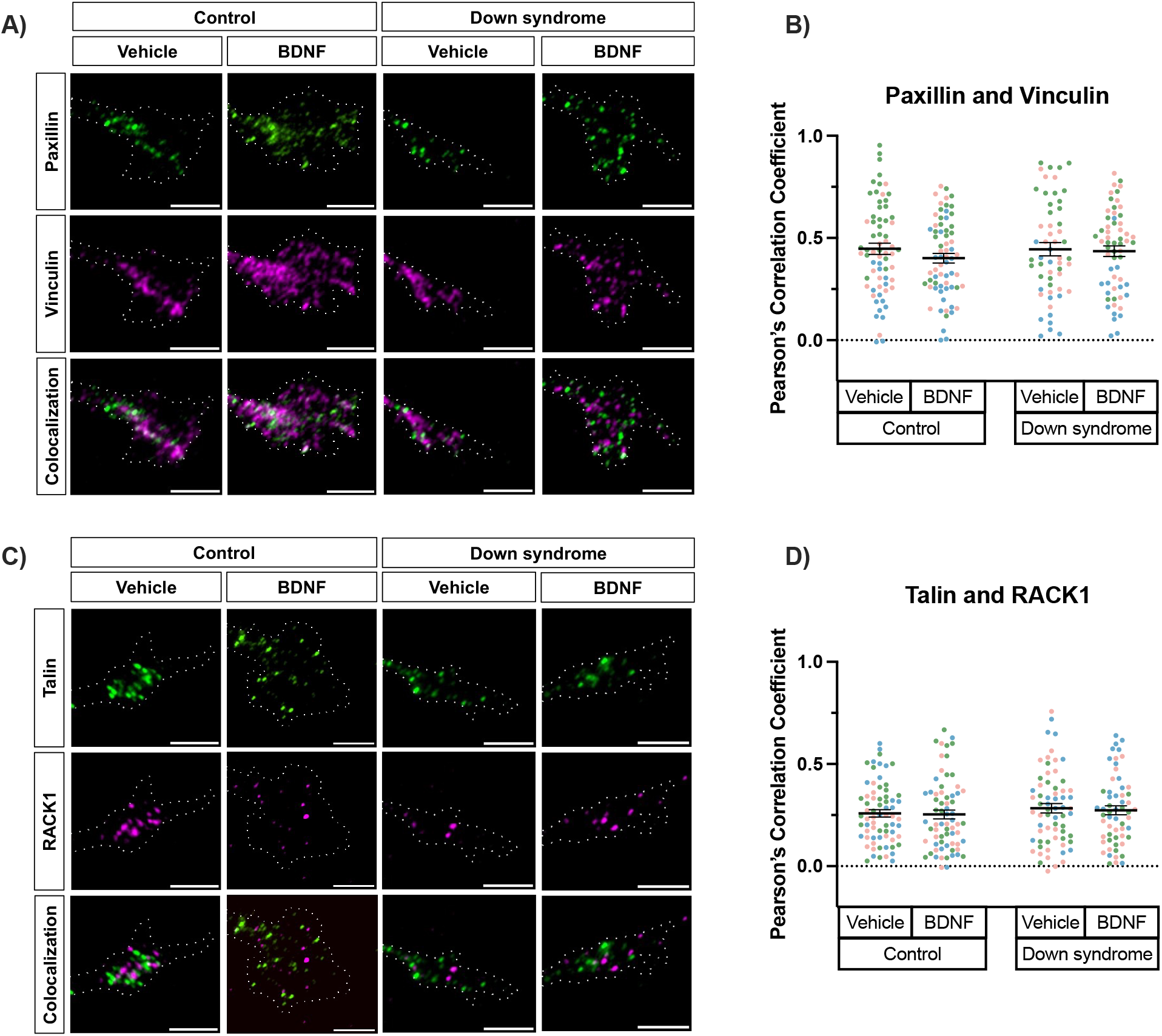
The colocalization of adhesion proteins in Down syndrome hiPSC-derived cortical neurons is not changed after BDNF stimulation. **(A, C)** Representative images of growth cones of control and Down syndrome hiPSC-derived cortical neurons that were stimulated with 100ng/ml BDNF or vehicle, and stained for adhesion proteins (paxillin, vinculin, talin, and RACK1) using quantitative immunofluorescence. The colocalization of paxillin and vinculin, and the colocalization of talin and RACK1 were quantified using Pearson’s correlation coefficient. Scale bars, 5μm. **(B)** Pearson’s correlation coefficient for paxillin and vinculin in growth cones of Control and Down syndrome hiPSC-derived cortical neurons under both vehicle and BDNF stimulated conditions. Two-way ANOVA: Genotype effect, p=0.5570; Treatment effect, p=0.3144; Interaction effect, p=0.4987. Control + Vehicle n=67; Control + BDNF n=67; Down syndrome + Vehicle n=54; Down syndrome + BDNF n=60. **(D)** Pearson’s correlation coefficient for talin and RACK1 in growth cones of Control and Down syndrome hiPSC-derived cortical neurons under both vehicle and BDNF stimulated conditions. Two-way ANOVA: Genotype effect, p=0.2812; Treatment effect, p=0.7174; Interaction effect, p=0.9175. Control + Vehicle n=69; Control + BDNF n=68; Down syndrome + Vehicle n=65; Down syndrome + BDNF n=58. “n” is the number of growth cones analyzed over 3 independent differentiations. Each differentiation set is signified by a separate color.

### Adhesion-associated genes display high inter-individual variability across multiple cell types in DS

We observed substantial inter-individual variability in adhesion protein expression in our dataset of T21 fibroblasts and iPSC-derived neurons. Using existing RNA-sequencing datasets, we compiled the RNA expression of the core adhesion proteins examined in this study (paxillin, vinculin, talin, and RACK1), as well as other critical adhesion members, including β-actin, focal adhesion kinase (FAK), laminins, and integrins, to determine whether this variability was widespread and if an adhesion component not studied herein showed more consistent changes in T21 (**Supplemental Table 3**). We included studies examining a range of cell types, including hiPSCs, hiPSC-derived neurons, human-derived neural progenitor cells (NPCs), fibroblasts, and postmortem cortical and hippocampal brain samples. We normalized the log2 fold-change (log2FC) expression levels between control and T21 cells to enable direct comparisons across T21 individuals and cell types (**Supplemental Table 3**).

To better visualize differential expression across genes in each study, the log2FC values were compiled into a heatmap (**Figure 8A**) and a bar graph highlighting the number of studies that reported the genes as upregulated, downregulated, or unchanged (**Figure 8B**). Paxillin (PXN) and talin (TLN1) were upregulated in half of the studies, while vinculin (VCL) showed more variable expression. Additionally, in all studies examining RACK1 (GNB2L1), it was not differentially expressed. Interestingly, other adhesion site components, such as alpha subunits of integrin receptors (ITGA1 and ITGA2) and subunits of the ECM protein laminin (LAMA1, LAMB1, and LAMC1), also showed variable gene expression, with 2-3 studies showing an increase and more showing no change. Together, these findings confirm that although adhesion-related genes are altered in Down syndrome, the specific proteins altered and the magnitude of the changes vary from individual to individual. By synthesizing multiple transcriptomic studies, we show that adhesion heterogeneity is reproducible across different studies and cell types.

**Figure 8.**
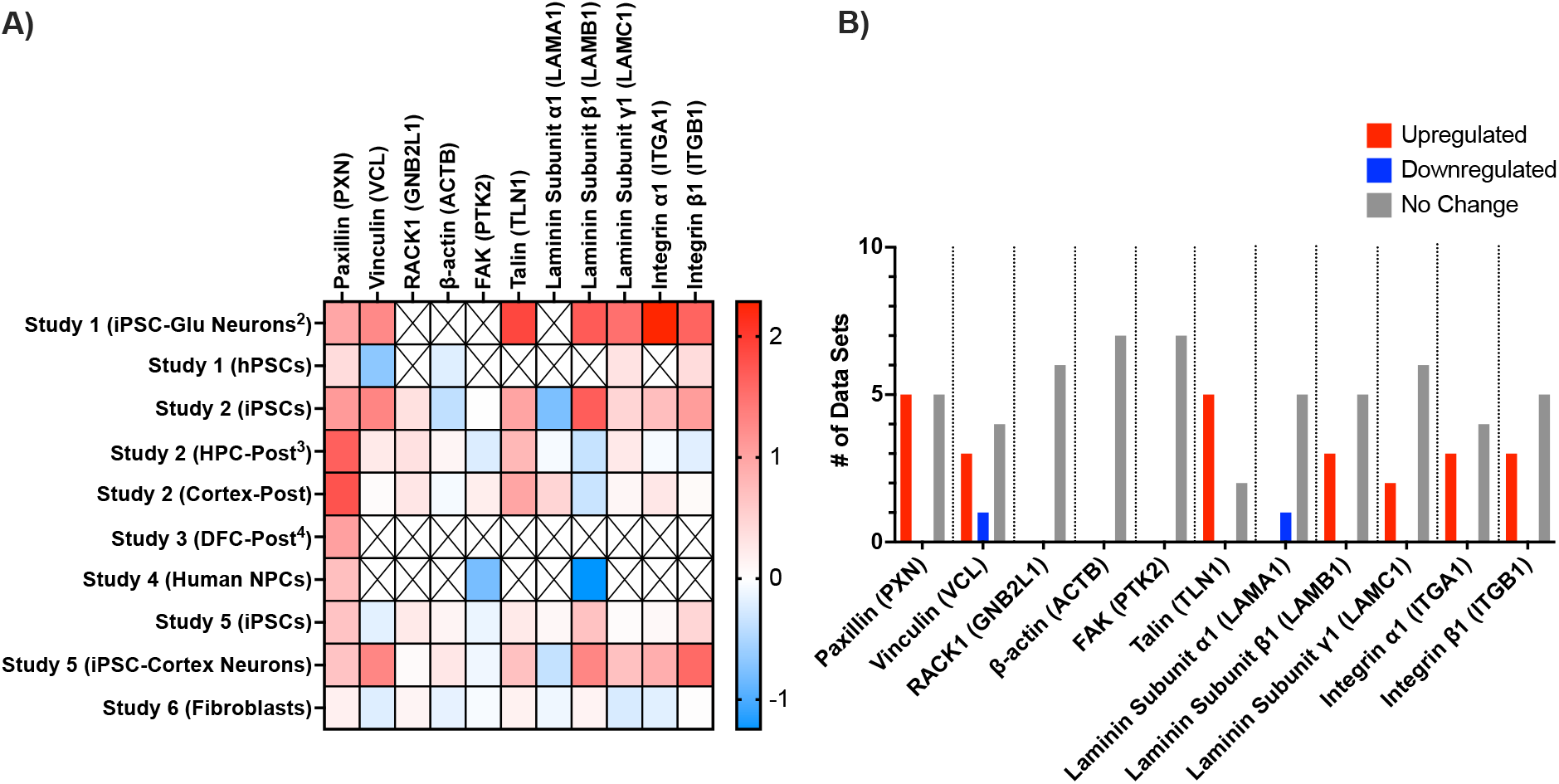
Differential regulation of adhesion and integrin-associated genes. **A)** Heatmap displaying log₂ fold-change (log₂FC) expression trends of adhesion proteins across multiple studies. **B)** Quantification of differential expression trends. The bar graph illustrates the number of data sets reporting upregulation (red), downregulation (blue), or no change (gray) for each gene of interest. Genes were considered significantly differentially expressed if they met the criteria of FDR (p-adjusted) < 0.05 and logFC > 0.58 or logFC < -0.58.

## DISCUSSION

Down syndrome results in many phenotypes, such as congenital heart disease, hypotonia, leukemia, slow wound healing, and intellectual disability (Antonarakis et al., 2020; Benvenuti et al., 2025; Bonnyman et al., 2024; Earley et al., 2022; Marentette et al., 2021; Murakami et al., 2008). However, the cellular and molecular mechanisms that regulate these phenotypes in Down syndrome remain poorly understood. We find altered cellular morphology and motility in both T21 fibroblasts and hiPSC-derived cortical neurons (**Figures 1, 2, and 4**), suggesting this may be a widespread cellular change contributing to multiple T21 phenotypes. Our previous work further supports this finding, as we have shown using both cortical neurons from the Ts65Dn mouse model of Down syndrome and hiPSC-derived cortical neurons that there are many changes in cellular morphology, including a decrease in axon length, soma area, branching, and growth cone area in T21 (Agrawal et al., 2026; Jain et al., 2020). Interestingly, we have also found that neuron motility is altered as axon guidance in response to the attractive guidance cue netrin-1 is lost in T21 hiPSC-derived cortical neurons (Agrawal et al., 2026).

The current findings are supported by previous studies that have reported changes in cellular morphology and motility in human iPSCs differentiated into multiple cell types. For example, T21 iPSC-derived GABAergic neurons are smaller, have fewer neurites, and exhibit reduced migration, both *in vitro* and *in vivo* after transplantation into the mouse medial septum (Huo et al., 2018). T21 iPSC-derived cranial neural crest cells and cardiomyocytes also exhibit significantly reduced migratory capacity (Liu et al., 2022; Reeser et al., 2024). Additionally, endothelial cells produced from T21 iPSCs have a decreased ability to form tubes (Moon and Lawrence, 2022; Perepitchka et al., 2020). Interestingly, iPSC-derived astrocytes also show changes, but in the opposite direction, with increased cell area and cellular motility (Ponroy Bally et al., 2020). This work on multiple iPSC-derived cell types is also supported by a study showing that microglial morphology is altered in post-mortem brain tissue from individuals with Down syndrome (Flores-Aguilar et al., 2020). Taken together with the current work, changes in cellular morphology and motility may be a ubiquitous phenotype of Down syndrome, irrespective of cell type; however, variations in the type and direction of changes occur both within and between cell types. It is also important to note that the identified changes in cellular morphology and motility occur across the lifespan. This is highly relevant to Down syndrome, as changes in cellular morphology and motility can contribute to dysregulated processes throughout life, for example, altered organ development, such as congenital heart disease, during early development, to changes in immune function in the adult.

Much remains to be understood about the molecular mechanisms underlying these cellular changes. Multiple sequencing studies have reported that grouping terms related to “adhesion” and “ECM” are some of the most dysregulated pathways in Down syndrome, with adhesion proteins more often upregulated than downregulated (**Figure 8**) (Conti et al., 2007; Guo et al., 2023; Huo et al., 2018; Meharena et al., 2022; Ponroy Bally et al., 2020; Rastogi et al., 2024; Sobol et al., 2019; Susco et al., 2022). For example, RNA-seq and ATAC-seq of iPSC-derived astrocytes found that extracellular matrix, cell-substrate junction, cell-substrate adherens junction, and focal adhesion are the processes with the greatest number of differentially expressed genes or accessible promoters in T21 (Ponroy Bally et al., 2020). Given that astrocytes secrete extensive ECM molecules in the brain, this work has implications for numerous nervous system cell types.

Some of the specific adhesion-related genes identified in these studies, and possibly leading to these changes, have been further investigated, but much remains poorly understood. Interestingly, the activation state of beta 1 integrin expression differs between T21 fibroblasts and leads to changes in adhesion (Jongewaard et al., 2002). Although this study found that beta 1 integrin expression is not significantly different between control and T21 fibroblasts (Jongewaard et al., 2002), our data comparing multiple RNA-seq studies show that it is upregulated in 3 out of 8 studies (**Figure 8**). Changes in specific adhesion and ECM-related proteins in Down syndrome have also been examined in the context of congenital heart defects (Mollo et al., 2023). Specifically, RUNX1, which is located on HSA21 and overexpressed in T21, is a transcription factor that regulates the expression of multiple ECM genes (Mollo et al., 2022). This study further showed that reducing RUNX1 expression could rescue reduced cellular migration. Another study has demonstrated that knockdown of the cell adhesion molecule CXADR can rescue migration defects present in T21 iPSC-derived cranial neural crest cells (Liu et al., 2022). Furthermore, a cell adhesion molecule, DSCAM, has also been more extensively studied in the context of Down syndrome, and reducing its levels has been shown to rescue multiple changes in T21, including neuritogenesis, synapse number, and axon guidance (Agrawal et al., 2026; Liu et al., 2023; Tang et al., 2021). Because a critical role of adhesions is to sense the extracellular matrix, changes in ECM composition may be an upstream driver of the changes identified herein. Two ECM proteins with strong evidence of changes in Down syndrome are collagens and fibronectin (Belgacemi et al., 2025; Conti et al., 2007; Galat et al., 2020; Gittenberger-de Groot et al., 2003; Karousou et al., 2013; Mollo et al., 2022; Mollo et al., 2023; Rodriguez-Lopez et al., 2024). We focused our studies on adhesion proteins rather than the ECM, but our studies suggest that the ECM should be the focus of future studies on T21, especially in the context of nervous system development and function, where there has been less focus.

We examined focal adhesions in fibroblasts, but found considerable variability between T21 cell lines (**Figure 3**). Interestingly, in the pairs that had decreased migration in the T21 line (Pair 1 and Pair 3), we also observed greater dysregulation of adhesion proteins (**Figure 3F**). When examining growth cones of T21 hiPSC-derived cortical neurons, the adhesion protein that showed the greatest change was RACK1 (**Figures 5E and 6E**), which is supported by another study showing that RACK1 is decreased in fetal brain from individuals with Down syndrome (Peyrl et al., 2002). RACK1 is a ribosomal binding protein that also localizes to and regulates adhesion sites (Buoso et al., 2022; Kershner and Welshhans, 2017). RACK1 binds to the classical ribonucleoprotein made up of β-actin mRNA and the RNA-binding protein, zipcode binding protein 1 (ZBP1) (Ceci et al., 2012). The local translation of β-actin, regulated by ZBP1 and RACK1, is necessary for cell motility and axon guidance (Ceci et al., 2012; Leung et al., 2006; Welshhans and Bassell, 2011; Yao et al., 2006). Thus, because RACK1 is a critical protein that regulates both adhesion signaling and translation, it is a compelling candidate to contribute to the changes described herein. The current data also underscore the importance of looking beyond the transcriptome, as RNA levels of RACK1 showed no change in any of the RNA-seq data sets examined (**Figure 8**), but we found changes at the protein level in half of the cell lines examined in this study.

Several limitations should be considered. Although multiple pairs of fibroblasts were examined, we only had access to a single set of human iPSCs, which limited our ability to assess neuronal inter-individual variability. Therefore, these observations require future validation in additional genetically independent lines. These experiments also established associations between some altered adhesion proteins and impaired cellular morphology, but did not directly test causality. In addition, we did not find differences in many of the adhesion proteins examined in this study; however, because we did not quantify these dynamic structures in live cells, we may have missed critical changes. Adhesion lifetime, turnover, assembly and disassembly rate, and traction force directly regulate cell morphology and motility (Mavrakis and Juanes, 2023). It is likely that measuring protein abundance alone may be insufficient to predict adhesion function because these are highly dynamic structures. Future work examining these adhesion proteins using live cell imaging is needed to further understand their contributions to the changes in cellular morphology and motility in T21. In addition, more research is needed on potential changes in mechanobiology in Down syndrome, which has been suggested by a recent study and could certainly be contributing to these cellular changes (Reeser et al., 2026). Future work should also examine whether altered adhesion dynamics arise through upstream signaling pathways such as Src/FAK.

These data also underscore the critical need in the field for clinical data. Given that we do see some adhesion proteins altered in a subset of individuals, such as integrins **(Figure 8**), it is critical to know if this subset also shares a similar phenotype, such as congenital heart disease or slow wound healing. With this information, we will be able to effectively integrate clinical data with cellular and molecular findings across large datasets, ultimately enabling the identification and investigation of molecular targets that may advance our understanding and treatment of Down syndrome. Interestingly, using data from hundreds of individuals, two recent studies have identified three main molecular subtypes of trisomy 21 and provided new insight into its clinical heterogeneity (Donovan et al., 2026; Donovan et al., 2024), which is a critical step in this direction and further supports the need for personalized medicine. These recently described molecular subtypes align with some of our data and thus provide a potential framework for explaining why distinct changes in adhesions can converge on similar cellular phenotypes.

We found high inter-individual variability in all parameters examined in this study. This could be due to a variety of reasons, including individuals’ genetic backgrounds, epigenetic differences, or differences in clinical profile. Thus, this work builds on the existing literature showing high inter-individual variation in Down syndrome, and a need for personalized medicine (Karmiloff-Smith et al., 2016; Lee et al., 2025; Upadhya et al., 2025; Waugh et al., 2025). The current work is significant because it goes beyond RNA-sequencing to determine whether changes in RNA are actually reflected at the protein level, as these often do not correlate (Srivastava et al., 2022; Vogel and Marcotte, 2012); this is demonstrated by the RACK1 data presented herein (**Figures 3, 5, 6, 8**). Overall, we do find high variability in adhesion proteins, and this is supported by the RNA-seq studies using hiPSCs, hiPSC-derived neurons, human- derived NPCs, fibroblasts, and postmortem cortical and hippocampal brain samples (**Supplemental Table 3; Figure 8**). Thus, the concordance between heterogeneous transcriptomic datasets and our data showing heterogeneous protein expression suggests that inter-individual variability is an intrinsic biological feature of Down syndrome.

In our study, we did not identify a single focal adhesion protein that may be responsible for the impaired morphology or motility in Down syndrome. Rather, our data, taken together with the broader literature, suggest that trisomy 21 alters an adhesion network at multiple nodes, in which distinct combinations of altered adhesion proteins, ECM molecules, and upstream pathways converge on common defects in cytoskeletal dynamics and cellular motility. In summary, this work demonstrates that there are robust, conserved cellular morphological and motility phenotypes of T21 that could be due, in part, to heterogeneous changes in adhesion.

Although phenotypes of Down syndrome, such as altered neural crest migration, wound healing, angiogenesis, and neural wiring, appear disparate, altered cellular morphology and motility fundamentally contribute to all of these. Further work is needed to identify the critical nodes underlying these changes, which can be used in personalized medicine to potentially treat phenotypes of Down syndrome that appear unrelated, but are united by shared changes in cellular structure.

## Acknowledgements

We thank Dr. Alberto Costa (Case Western University; Cleveland, Ohio) for the control and Down syndrome hiPSCs.

## FIGURES & FIGURE LEGENDS

**Supplemental Table 1.**
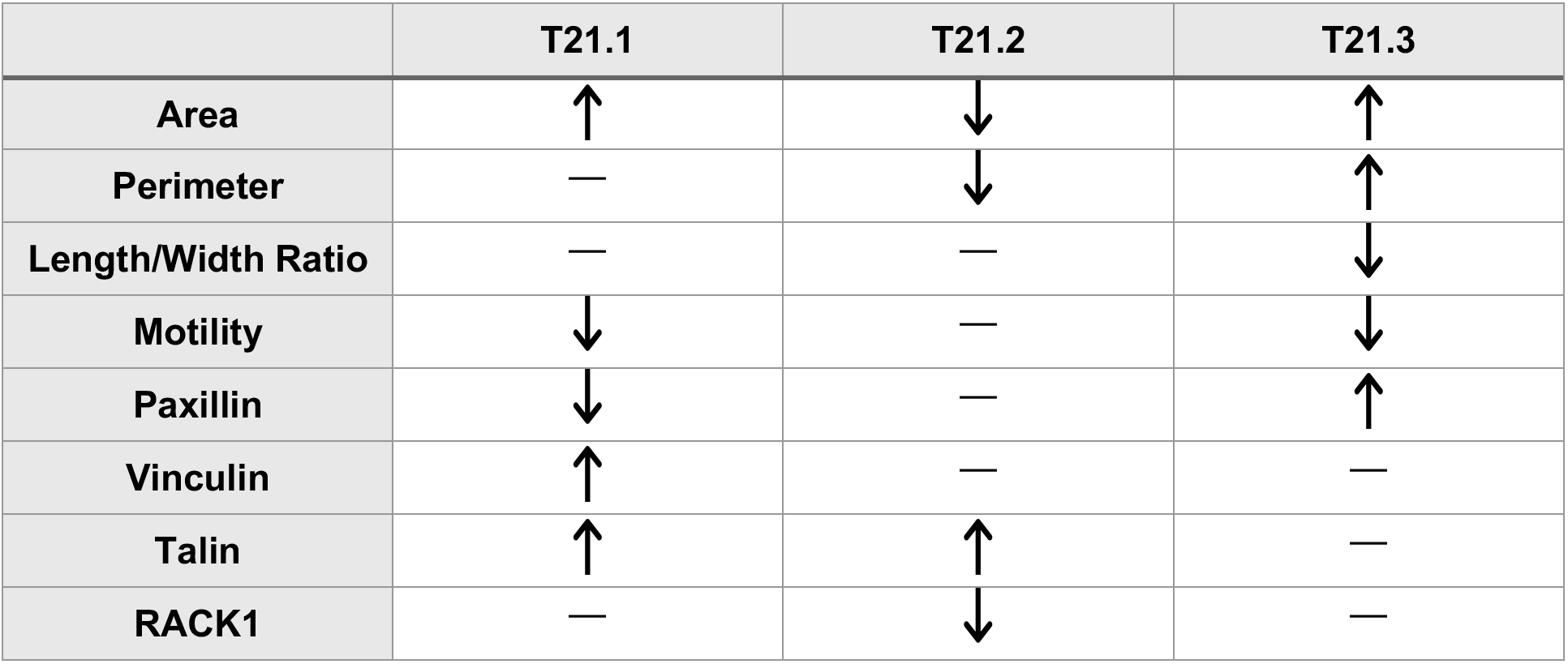
Summary of changes in Down syndrome fibroblast morphology, motility, and protein expression.

|  | T21.1 | T21.2 | T21.3 |
| --- | --- | --- | --- |
| **Area** | ↑ | ↓ | ↑ |
| **Perimeter** | — | ↓ | ↑ |
| **Length/Width Ratio** | — | — | ↓ |
| **Motility** | ↓ | — | ↓ |
| **Paxillin** | ↓ | — | ↑ |
| **Vinculin** | ↑ | — | — |
| **Talin** | ↑ | ↑ | — |
| **RACK1** | — | ↓ | — |

**Supplemental Table 2.** Summary of changes in morphology, adhesion protein expression, and colocalization of adhesion proteins in growth cones of T21 iPSC-derived neurons compared to control iPSC-derived neurons for each treatment condition (i.e., at basal conditions, starved conditions, and following BDNF stimulation).

|  | T21 Neurons - Basal | T21 Neurons - Starved | T21 Neurons - BDNF |
| --- | --- | --- | --- |
| **Growth Cone Area** | ↓ | N/A | N/A |
| **Longest Neurite Length** | ↓ | N/A | N/A |
| **Paxillin** | — | — | — |
| **Vinculin** | — | — | — |
| **Talin** | ↑ | — | — |
| **RACK1** | ↑ | — | ↑ |
| **Paxillin + Vinculin** | N/A | — | — |
| **Talin + RACK1** | N/A | — | — |

**Supplemental Table 3.** Normalized Log_2_ Fold-Change (Log_2_FC) values of adhesion and integrin-associated proteins across studies.

| Study | Reference | Cell Type | PXN | VCL | GNB2L1 | ACTB | PTK2 | TLN1 | LAMA1 | LAMB1 | LAMC1 | ITGA1 | ITGB1 |
| --- | --- | --- | --- | --- | --- | --- | --- | --- | --- | --- | --- | --- | --- |
| Study 1 | (Susco et al., 2022) | iPSC-Glu Neurons<sup>2</sup> | 0.973* | 1.27* | N/A | N/A | N/A | 1.89* | N/A | 1.71* | 1.51* | 2.29* | 1.62* |
|  |  | hPSCs | 0.396 | -0.685* | N/A | -0.207 | N/A | N/A | N/A | N/A | 0.317 | N/A | 0.396 |
| Study 2 | (Rastogi et al., 2024) | iPSCs | 1.09* | 1.31* | 0.343 | -0.391 | 0.00778 | 1.002* | -0.769* | 1.68* | 0.471 | 0.722* | 1.07* |
|  |  | HPC-Post<sup>3</sup> | 1.66* | 0.239 | 0.339 | 0.109 | -0.228 | 0.794* | -0.0589 | -0.358 | 0.245 | -0.0609 | -0.197 |
|  |  | Cortex-Post | 1.77* | 0.0427 | 0.285 | -0.0636 | 0.196 | 0.989* | 0.476 | -0.338 | 0.0953 | 0.273 | 0.0625 |
| Study 3 | (Olmos-Serrano et al., 2016) | DFC-Post<sup>4</sup> | 1.03* | N/A | N/A | N/A | N/A | N/A | N/A | N/A | N/A | N/A | N/A |
| Study 4 | (Esposito et al., 2008) | Human NPCs | 0.705 | N/A | N/A | N/A | -0.791 | N/A | N/A | -1.25 | N/A | N/A | N/A |
| Study 5 | (Gonzales et al., 2018) | iPSCs | 0.66 | -0.18 | 0.24 | 0.12 | -0.13 | 0.24 | 0.09 | 0.68 | 0.04 | 0.08 | 0.46 |
|  |  | iPSC-Cortex Neurons | 0.66 | 1.3* | 0.05 | 0.27 | -0.1 | 0.69* | -0.37 | 1.3* | 0.69* | 0.89* | 1.57* |
| Study 6 | (Sullivan et al., 2016) | Fibroblasts | 0.166 | -0.208 | 0.123 | -0.158 | -0.0547 | 0.151 | -0.107 | 0.137 | -0.249 | -0.194 | 0.0156 |

\* – Indicates statistically significant difference
<sup>1</sup> N/A – Protein was not detected in the study or was not significantly different ( $p > 0.05$ ).
<sup>2</sup> iPSC-Glu Neurons – iPSC-derived Glutamatergic Neurons.
<sup>3</sup> HPC-Post – Hippocampus postmortem samples.
<sup>4</sup> DFC-Post – Dorsal frontal cortex postmortem samples.

